# Microbial regulation of hexokinase 2 links mitochondrial metabolism and cell death in colitis

**DOI:** 10.1101/2020.12.22.423953

**Authors:** Jacob Hamm, Finn Hinrichsen, Lena Schröder, Neha Mishra, Kensuke Shima, Alesia Walker, Nina Sommer, Kenneth Klischies, Daniela Prasse, Johannes Zimmermann, Sina Kaiser, Dora Bordoni, Antonella Fazio, Georg Laue, Valentina Tremaroli, Marijana Basic, Robert Häsler, Ruth A. Schmitz, Stefan Krautwald, Andrea Wolf, Bärbel Stecher, Philippe Schmitt-Kopplin, Christoph Kaleta, Jan Rupp, Fredrik Bäckhed, Philip Rosenstiel, Felix Sommer

## Abstract

Hexokinases (HK) catalyze the first step of glycolysis and thereby limit its pace. HK2 is highly expressed in the gut epithelium, plays a role in immune responses and is upregulated in inflammation and ulcerative colitis ^1–3^. Here, we examined the microbial regulation of HK2 and its impact on intestinal inflammation by generating mice lacking HK2 specifically in intestinal epithelial cells (*Hk2*^Δ*IEC*^). *Hk2*^Δ*IEC*^ mice were less susceptible to acute intestinal inflammation upon challenge with dextran sodium sulfate (DSS). Analyzing the epithelial transcriptome from *Hk2*^Δ*IEC*^ mice during acute colitis revealed downregulation of cell death signaling and mitochondrial dysfunction dependent on loss of HK2. Using intestinal organoids derived from *Hk2*^Δ*IEC*^ mice and Caco-2 cells lacking HK2, we identified peptidyl-prolyl cis-trans isomerase (PPIF) as a key target of HK2-mediated regulation of mitochondrial permeability and repression of cell-death during intestinal inflammation. The microbiota strongly regulated HK2 expression and activity. The microbially-derived short-chain fatty acid (SCFA) butyrate repressed HK2 expression and oral supplementation protected wildtype but not *Hk2*^Δ*IEC*^ mice from DSS colitis. Our findings define a novel mechanism how butyrate may act as a protective factor for intestinal barrier homeostasis and suggest targeted HK2 inhibition as a promising therapeutic avenue in intestinal inflammation.

---

Hexokinases (HK) catalyze the first step of glycolysis and thereby limit the rate of this fundamental biological process. HK2 is considered the prototypic inducible isoform of all HK family members as it can be upregulated by various environmental factors and signaling pathways, e.g. during inflammation and in ulcerative colitis ^1–3^. In addition to its metabolic function, HK2 acts as a receptor for bacterial cell wall components ^4^ and has been suggested to counteract mitochondria-mediated cell death ^5^. Chemical inhibition of HK impairs immune cell activation and promotes infection by *Listeria monocytogenes* in immune cells ^6^, whereas hyperglycemia and high glycolytic flux have been associated with an increased risk of enteric infection in intestinal epithelial cells ^7^. However, complete ablation of HK2 is embryonically lethal ^8^. In the intestine, *Hk2* is predominantly expressed by intestinal epithelial cells (IECs) ^9,10^. Therefore, we aimed to investigate, whether selective ablation of HK2 in IECs alters epithelial function during intestinal inflammation. Here we show that (i) loss of HK2 in IECs protects from DSS-induced colitis by decreasing inflammation-induced epithelial cell death and that (ii) specific bacterial species and the microbial metabolite butyrate ameliorate colitis by repressing *Hk2* expression.

To determine the role of epithelial HK2 for intestinal inflammation, we generated Hk2^fl/fl^-Villin:: *Cre*^+^ mice lacking HK2 specifically in IECs, hereinafter referred to as *Hk2*^Δ*IEC*^ mice. We used littermate Hk2^fl/fl^ mice, hereinafter referred to as WT mice, as controls. Unchallenged *Hk2*^Δ*IEC*^ mice did not display any major inflammatory or metabolic phenotype except for an improved glucose tolerance (Extended data Figure 1), compared to littermate controls. However, when *Hk2*^Δ*IEC*^ mice and their WT littermates were challenged with dextran sodium sulfate (DSS) to induce intestinal inflammation, *Hk2*^Δ*IEC*^ mice lost significantly less weight compared to WT littermates (Fig. 1a). The disease activity index (DAI), a measure of intestinal inflammation comprised of weight loss, stool consistency and fecal blood occurrence, confirmed the ameliorated disease course in *Hk2*^Δ*IEC*^ mice (Fig. 1b). Additionally, *Hk2*^Δ*IEC*^ mice displayed lower serum levels of the pro-inflammatory cytokine KC/CXCL1 as measured by ELISA (Fig. 1c). Histological evaluation of Hematoxylin and Eosin (H&E)-stained colon sections demonstrated a reduced score consisting of transmural inflammation, crypt hyperplasia, epithelial injury, polymorphonuclear and mononuclear cell infiltrates in *Hk2*^Δ*IEC*^ mice (Fig. 1d). In WT mice, HK2 expression increased during the course of colitis (Fig. 1e, Extended data figure 2). We therefore investigated whether *Hk2* expression is dysregulated in patients suffering from intestinal inflammation by evaluating expression data from colon-biopsies of patients suffering from Crohn’s Disease (CD), Ulcerative Colitis (UC) or Non-IBD-Colitis (NIC) ^11^. Biopsies were taken both from the site of inflammation and adjacent non-inflamed tissue. We found that *Hk2* expression was significantly higher in the inflamed tissue of CD (p = 0.010) and NIC (p = 0.027) patients, while only a trend was observed in UC patients (p = 0.180) (Fig. 1f). This data is in agreement with a published report revealing upregulation of *Hk2* expression in UC patients suggesting inhibition of HK2 as a potential therapeutic approach ^3^. To decipher the molecular mechanisms protecting *Hk2*^Δ*IEC*^ mice from intestinal inflammation, we isolated IECs from WT and *Hk2*^Δ*IEC*^ mice on day 0 (baseline), 3 (early inflammation) and 7 (late inflammation) from an independent acute DSS experiment and performed RNA sequencing. While we did not find any differentially expressed genes on day 3, we identified 420 differentially expressed genes on day 7 in comparison between WT and *Hk2*^Δ*IEC*^ mice (Extended data Figure 3 and Extended data Table 1). Gene Ontology (GO) analysis of these differentially expressed genes revealed a downregulation of genes involved in cell death-signaling and regulation of mitochondrial membrane permeability in *Hk2*^Δ*IEC*^ mice (Fig. 1g). We confirmed a reduction of cell death in *Hk2*^Δ*IEC*^ mice by TUNEL-staining of colon sections from day 3 and day 7 of the experiment. In accordance with our transcriptomics data, we found fewer TUNEL-positive cells in the tip compartment of colonic crypts in *Hk2*^Δ*IEC*^ mice at day 7 indicating less cell death (Fig. 1h). This finding coincides with the spatial expression profile of HK2, which is mainly restricted to the colonic epithelial tip (Extended data Figure 4).

**Figure 1:**
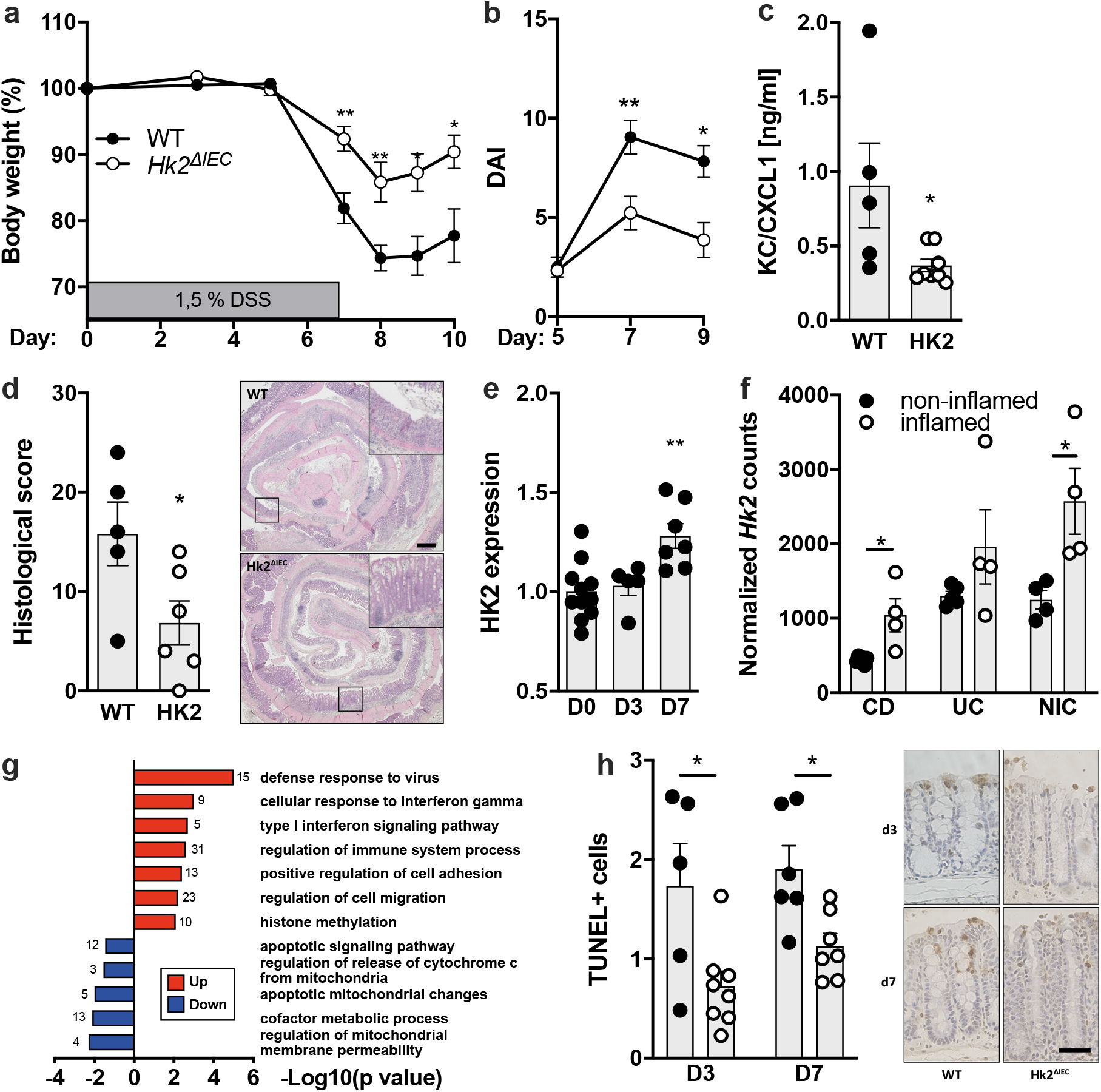
Loss of HK2 in the intestinal epithelium protects from colitis. **a)** Body weight loss of WT and *Hk2*^ΔIEC^ mice lacking HK2 in intestinal epithelial cells during DSS-induced colitis. * p<0.01 and ** p<0.01 WT versus *Hk2*^ΔIEC^ mice using two-way ANOVA. **b)** Disease activity index (DAI) consisting of stool consistency, fecal blood occurrence and body weight loss. * p<0.01 and ** p<0.01 WT versus *Hk2*^ΔIEC^ mice using two-way ANOVA. **c)** KC/CXCL1 (pro-inflammatory cytokine) levels in serum of WT and *Hk2*^ΔIEC^ mice as determined by ELISA. * p<0.01 using Mann-Whitney U-test. **d)** Histological score of H&E-stained sections from colon of WT and *Hk2*^ΔIEC^ mice including representative images of the experimental groups. The scale bar represents 500 μm. * p<0.01 using Mann-Whitney U-test. **e)** Relative HK2 protein expression during the course of DSS-induced colitis in colon epithelium of WT mice as determined by immunohistochemistry. ** p<0.01 versus D0 using one-way ANOVA. **f)** *Hk2* expression is dysregulated in inflamed human intestinal mucosa. *Hk2* expression was determined in inflamed and non-inflamed intestinal mucosal biopsies from patients with Crohn’s Disease (CD), Ulcerative Colitis (UC) or non-IBD Colitis (NIC) by RNA-Seq. n=4-6 per group. * p<0.05. **g)** Gene ontology terms enriched in up- and down-regulated genes in transcriptomes of colonic IEC isolated from *Hk2*^ΔIEC^ compared to WT mice sacrificed on day 7 of DSS colitis. **h)** Fewer apoptotic cells per colon crypt in *Hk2*^ΔIEC^ mice as determined by TUNEL assay including representative images. The scale bar represents 50 μm. * p<0.01 using two-way ANOVA.

To further study the molecular processes involved in HK2-dependent protection from inflammation, we generated intestinal organoids derived from *Hk2*^Δ*IEC*^ and WT mice and investigated their response to stimulation with tumor necrosis factor (TNF). Western blot analysis demonstrated higher HK2 levels upon TNF stimulation (Fig. 2a). Organoids derived from *Hk2*^Δ*IEC*^ mice exhibited lower levels of cleaved Caspase 3 and Poly(ADP-Ribose)-Polymerase 1 (PARP1), markers for mitochondria-related types of cell death, compared to WT organoids upon TNF stimulation. This data therefore supported reduced levels of cell death in the absence of HK2 under inflammatory conditions. Furthermore, our transcriptome data suggested dysregulated mitochondrial function as a conse-quence of loss of HK2. We therefore performed metabolic flux analysis using Seahorse technology. As the 3D structure of organoids limits their use in this assay, we generated a Caco-2 cell clone lacking HK2 using the CRISPR Cas9 system, hereinafter named *Caco-2*^Δ*Hk2*^. We assessed glycolytic flux by measuring the extracellular acidification rate (ECAR) upon addition of glucose to induce glycolytic flux, oligomycin to stress the glycolytic reserve and 2-desoxy-glucose to inhibit glycolysis. Throughout the entire experiment, we did not observe any significant changes between Caco-2^Δ*Hk2*^ and Caco-2^WT^ cells indicating that ablation of HK2 did not affect glycolytic function (Fig. 2b). We also measured the basal oxygen consumption rate (OCR), which comprises both mitochondrial and non-mitochondrial oxygen consumption. Interestingly, we discovered significantly lower basal mitochondrial respiration as well as a dras-tically lower maximal mitochondrial respiration in *Caco-2*^Δ*Hk2*^ cells compared to Caco-2^WT^ cells (Fig. 2c). Since FCCP (cyanide-4-(trifluoromethoxy)-phenylhy-drazone) induces maximal respiration by depolarization of the mitochondrial membrane, our data therefore pointed towards a decrease in mitochondrial permeability. These results support our findings from transcriptome sequencing and GO analysis, which also indicated a decreased regulation of mitochondrial permeability. We therefore screened our transcriptome data for differentially expressed genes that are involved in regulation of mitochondrial membrane permeability. mRNA levels of *Ppif* (peptidyl-prolyl cis-trans isomerase), encoding for a main component of the mitochondrial permeability transition pore (MPTP), were downregulated in IECs of Hk2^Δ*IEC*^ mice on day 7 of colitis (Fig. 2d). PPIF coordinates mitochondrial permeability and metabolism ^12–15^ and has been suggested to directly interact with HK2 to suppress cell death ^5^. To validate our *in vivo* findings and transcriptome data, we stimulated *Caco-2*^Δ*Hk2*^ and Caco-2^WT^ cells with TNF and IL17A ^16^ as well as with IFN-β to induce inflammatory responses and cell death. Indeed, *Ppif* expression was downregulated under both conditions (Fig. 2e). Based on our findings and since PPIF interacts with HK2 ^5^ and since *Ppif*^-/-^ mice are less susceptible to colitis ^17^, we propose that mechanistically the HK2-dependent protection from intestinal inflammation could be mediated by lower levels of PPIF and a subsequent decrease in MPTP opening and mitochondrial membrane permeability.

**Figure 2:**
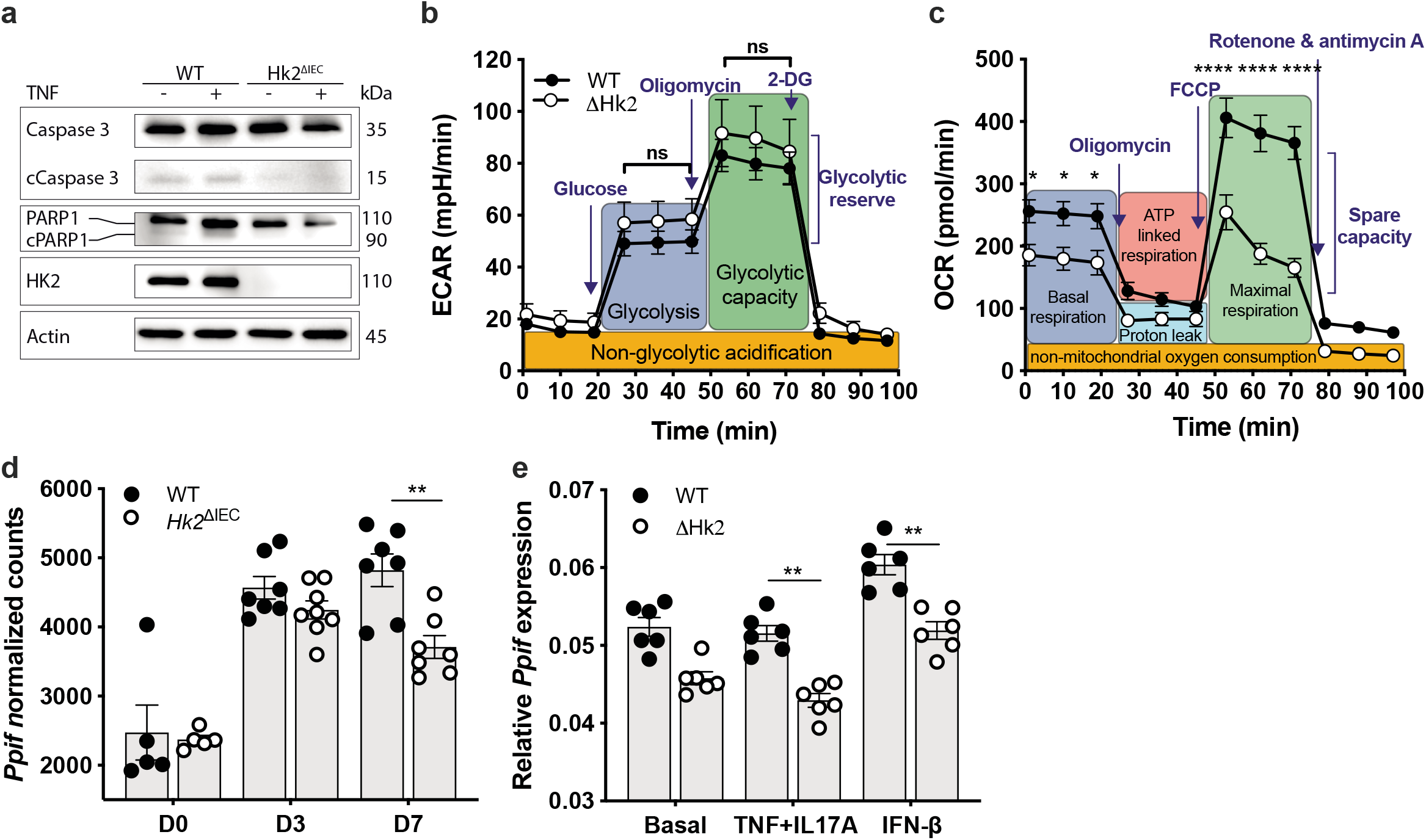
Dysregulated mitochondrial function in response to loss of HK2. **a)** Western blot analysis protein lysates of intestinal organoids raised from WT and *Hk2*^ΔIEC^ mice after stimulation with 100 ng/ml TNF for 24h. **b,c)** Metabolic analysis of WT and *Hk2*-deficient Δ*Hk2* Caco-2 cells using the Seahorse XF analyzer. * p<0.01 and **** p<0.0001 WT versus Δ*Hk2* using two-way ANOVA. **b)** The extracellular acidification rate (ECAR) reflects the glycolytic flux (ns = non-significant). **c)** The oxygen consumption rate (OCR) indicates mitochondrial respiration, which was impaired due to loss of HK2. * p<0.05, **** p<0.0001. **d)** Epithelial *Ppif* expression was downregulated in *Hk2*^ΔIEC^ compared to WT mice during the course of DSS-induced colitis (days 0, 3 and 7) as determined by RNA sequencing (normalized read counts). ** p<0.01 WT versus *Hk2*^ΔIEC^ using two-way ANOVA. **e)** Reduced *Ppif* expression as measured by qPCR in Δ*Hk2* Caco-2 compared to WT cells upon 24h stimulation with TNF (100 ng/ml) and IL17A (50 ng/ml) or with IFN-β (50 ng/ml) to induce inflammation. ** p<0.01 WT versus Δ*Hk2* using two-way ANOVA.

Dysregulated host-microbiota interactions are a key element of intestinal in-flammation ^18^. By comparing the transcriptomes of intestinal epithelial cell fractions isolated from germ-free (GF) and conventionally raised (CR) mice ^19^, we identified HK2 as significantly upregulated by the presence of a complex microbial community. *Hk2* expression was specifically induced in the epithelial tips of both ileum and colon by the microbiota (Fig. 3a). Upon colonization of GF mice with a normal microbiota *Hk2* expression increased and normalized to that of CR mice, demonstrating that the intestinal microbiota stimulates *Hk2* expression (Fig. 3b). However, although we were able to show a distinct effect of the microbiota on *Hk2* expression, ablation of HK2 in IECs in turn did not impact the composition of the intestinal microbiota as assessed by 16S rRNA amplicon sequencing under basal unchallenged conditions (Extended data Fig. 5). Next, we investigated how the microbiota regulates epithelial *Hk2* expression, specifically whether only specific bacterial species modulate *Hk2* expression. To that end, we colonized GF mice with either single bacterial species or minimal consortia - namely the Altered Schaedler Flora (ASF) ^20^ and the Oligo Mouse Microbiota (OMM) ^21^. Both minimal microbial consortia readily induced *Hk2* expression to a similar level as observed in CR mice (Fig. 3c). Mono-colonization with the Gram-negative bacterium *Bacteroides thetaiotaomicron* was also able to induce *Hk2* expression, whereas the Gram-negative bacterium *Escherichia coli* and the Gram-positive bacterium *Bifidobacterium longum* did not alter *Hk2* mRNA levels (Fig. 3c). Together, this data suggested a specific interaction between specific bacterial features and epithelial cells rather than general principles such as recognition of lipopolysaccharide or peptidoglycan as a mechanism regulating HK2 expression. We thus next aimed to disentangle the effects of individual OMM bacteria to identify potential candidate principles regulating *Hk2* expression. To that end, we stimulated Caco-2 cells with sterile-filtered culture supernatants of the individual species of the OMM consortium. We identified *Enterococcus faecalis* KB1 as the key inducer among this minimal microbiota (Fig. 3d). *Clostridium innocuum* I46 and *Flavonifractor plautii* YL31, both well-known producers of short-chain fatty acids (SCFA) ^22^, significantly reduced *Hk2* expression (Fig. 3d). As other fatty acids such as palmitic acid can inhibit HK activity ^23,24^, we hypothesized that SCFAs could drive the regulation of *Hk2* expression. To test this hypothesis, we generated individual metabolic models predicting the SCFA synthesis potential for each OMM member (Extended data Figure 6a). A linear model of the predicted *Hk2* expression based on the butyrate and acetate levels significantly correlated with the experimental *Hk2* expression determined by qPCR (p value = 0.008, Pearson R = 0.75, R^2^ = 0.56, AIC = −8.0; Extended data Figure 6b and Extended data Table 2). The predicted SCFA levels in the OMM culture supernatants were validated by targeted metabolomics (Extended data Figure 6c). We next stimulated Caco-2 cells with butyrate and acetate and found that acetate upregulated, whereas butyrate downregulated *Hk2* expression (Fig. 3e). We then tested whether stimulation of Caco-2 cells with butyrate also affected *Ppif* expression. Indeed, butyrate also downregulated *Ppif* expression (Fig. 3f). Together this data suggested that the microbial metabolite butyrate could potentially protect from inflammation via HK2- and PPIF-mediated changes in mitochondrial function and cell death. We therefore tested whether dietary supplementation of butyrate also functions *in vivo* to downregulate HK2 levels. We fed a butyrate-enriched diet to WT mice for 10 days and quantified HK2 levels in colon sections by immunohistochemistry, which demonstrated a clear downregulation of HK2 (Fig. 3g). We then set out to test whether the SCFA-dependent modulation of HK2 levels also impacts colitis outcome. Therefore, we supplemented *Hk2*^Δ*IEC*^ mice and their WT littermates with three different diets – a control diet, a butyrate-enriched diet or an acetate-containing diet – and induced colitis by performing the DSS colitis model as before. Indeed, in WT mice dietary supplementation of butyrate ameliorated colitis, which was evident by less weight loss (Fig. 3h), a lower DAI (Fig. 3i), lower serum levels of KC/CXCL1 (Fig. 3j), fewer TUNEL-positive epithelial cells (Fig. 3k) and a lower histological score (Fig. 3l). In contrast, acetate supplementation worsened colitis outcome as evident by increased weight loss (Fig. 3h) and a higher DAI (Fig. 3i), which even led to premature termination of this experimental group for ethical reasons on day 8. Acetate supplementation significantly increased whereas butyrate lowered colonic HK2 levels in WT mice (Fig. 3m). In *Hk2*^Δ*IEC*^ mice, treatment with butyrate did not impact colitis outcome as measured by weight development, DAI, serum KC/CXCL1 levels, TUNEL-positive epithelial cells or histological score (Fig. 3 h-l and Extended data figure 7). Together this data therefore demonstrates that ablation of HK2 in the intestinal epithelium completely blunted the butyrate-dependent effects on colitis. SCFAs are well-known for their pleiotropic effects on host physiology including intestinal motility, inflammation and carcinogenesis ^25^. Regarding colitis, acetate was shown to exacerbate ^26^ whereas butyrate ameliorates ^27–29^ inflammation. Butyrate is produced by the metabolic activity of the colonic microorganisms through fermentation of dietary fiber ^30^. Mechanistically, butyrate is sensed by the G protein-coupled receptors GPR41 (FFAR3) ^31,32^, GPR43 (FFAR2) ^31,33–35^ and GPR109a (HCA2) ^29,36^, which could trigger a signaling response leading to reduced *Hk2* expression. However, silencing these GPRs by siRNAs did not alter repression of *Hk2* expression by butyrate in Caco-2 cells (Extended data Figure 6d-e), which argues against a prominent role of these GPRs. Alternatively, butyrate also directly acts on histone deacetylases (HDACs) ^25^, which function as epigenetic regulators and thereby could impact on *Hk2* expression. Using either a pan-HDAC inhibitor or those specific for single HDAC classes or enzymes, we found that class I HDACs, possibly HDAC2 or HDAC3 mediate the repression of *Hk2* expression by butyrate (Extended data Figure 6f).

**Figure 3:**
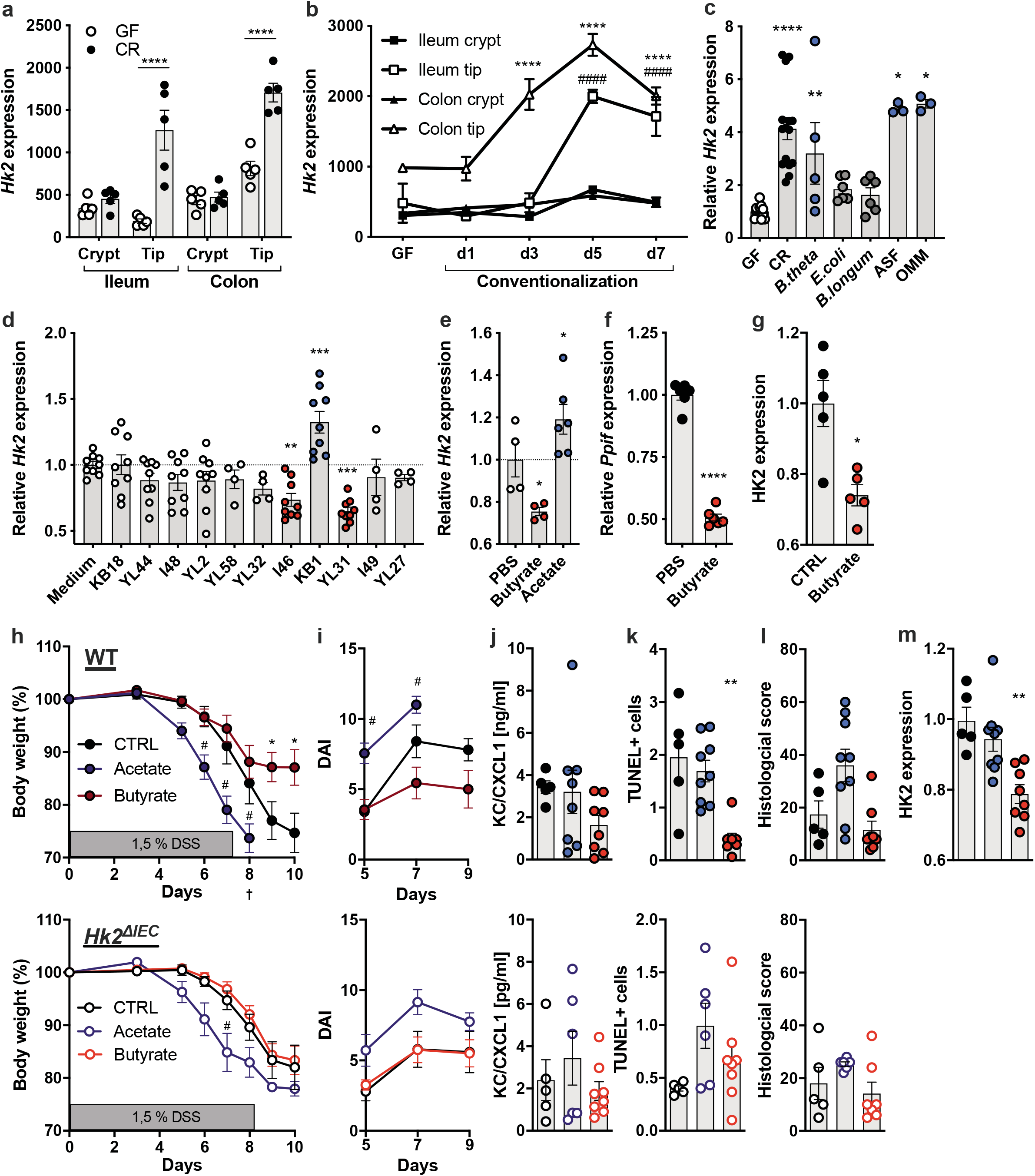
The microbial butyrate ameliorates colitis via HK2. *Hk2* expression in crypts and tips of ileum and colon of **a)** germ-free (GF) and conventionally raised (CR) mice and **b)** during colonization of GF mice with a normal microbiota. **** p<0.0001 GF vs. CR using moderated t test with FDR correction. # p<0.001 GF vs. d1 or d3 or d5 or d7 using one-way ANOVA. **c)** *Hk2* expression in GF mice and those mono-colonized with the single bacteria *Bacteroides thetaiotaomicron* (Gram-), *Escherichia coli* (Gram-) or *Bifidobacterium longum* (Gram+) or the minimal microbiomes ASF (Altered Schaedler Flora) and OMM (Oligo-Mouse-Microbiota). * p<0.01, ** p<0.01 and **** p<0.0001 versus GF using one-way ANOVA. **d)** Relative *Hk2* expression in Caco-2 cells stimulated with sterile-filtered culture supernatants of the OMM species grown *in vitro*. The used strains were: *Acutalibacter muris* KB18*, Akkermansia municiphila* YL44*, Bacteroides ceacimuris* I48*, Bifidobacterium animalis* YL2*, Blautia coccoides* YL58*, Enterocloster clostridioforme YL32, Clostridium innocuum* I46, *Enterococcus faecalis* KB1*, Flavonifractor plautii* YL31*, Limosilactobacillus reuteri* I49*, Muribaculum intestinale* YL27. ** p<0.01 and *** p<0.001 versus Medium using one-way ANOVA. **e)** Relative *Hk2* expression in Caco-2 cells stimulated with the microbial metabolites acetate and butyrate (10 mM each for 24h). * p<0.01 versus PBS using one-way ANOVA. **f)** Relative *Ppif* expression in Caco-2 cells stimulated with butyrate. **** p<0.0001 versus PBS using one-way ANOVA. **g)** Feeding WT mice a butyrate-enriched diet reduced HK2 protein expression in colonic epithelium as measured by immunohistochemistry. * p<0.01 versus CTRL diet using one-way ANOVA. **h-m)** Dietary supplementation of the microbial metabolite butyrate protected from colitis dependent on HK2. WT and *Hk2*^ΔIEC^ mice were fed either an acetate-enriched, butyrate-enriched or control diet and were then orally administered DSS to induce colitis. **h)** Body weight development, **i)** DAI, **j)** serum KC/CXCL1 levels, **k)** TUNEL-positive cells per colon crypt, **l)** histological score of WT (upper panel) and *Hk2*^ΔIEC^ (lower panel) mice. **m)** HK2 protein levels in colon epithelium of treated WT mice as per immunohistochemistry. Acetate stimulated whereas butyrate repressed HK2 expression. * / # indicate statistical significance in Butyrate (*) or Acetate (#) versus CTRL diet using two-way ANOVA in (**h-i**) and one-way ANOVA in (**j-m**).

Taken together, our study revealed a novel regulatory circuit consisting of epi-thelial HK2 and the microbial metabolite butyrate. We found that ablation of HK2 protects from colitis by suppression of cell death that was linked to altered mitochondrial function, which could be due to PPIF-dependent opening of the MPTP. Moreover, we identified the intestinal microbiota and its metabolite bu-tyrate as potent regulators of HK2 and we demonstrated that the protective effect of dietary butyrate supplementation is dependent on the functional presence of HK2. Previous studies already pointed towards a beneficial function of butyrate-producing bacteria for a healthy intestine and prevention of gut inflammation ^37–42^ and clinical trials using oral supplementation of germinated barley foods that are fermented into SCFA by the microbiota or rectal enema with butyrate indeed demonstrated beneficial effects in ulcerative colitis patients ^43,44^. Our findings therefore shed light on the molecular mechanism how butyrate mediates its beneficial effects and may guide the development of more specific therapeutic options by targeting HK2.

## Methods

### Animals

All animal experiments were approved by the local animal safety review board of the federal ministry of Schleswig Holstein and conducted according to national and international laws and policies (V 312-72241.121-33 (95-8/11) and V242-62324/2016 (97-8/16)). Specific-pathogen free (SPF) animals were housed in the Central Animal Facility (ZTH) of the University Hospital Schleswig Holstein (UKSH, Kiel, Germany). To create the Hk2^ΔIEC^ mouse line, we crossed commercially available mice carrying a floxed *Hk2* allele (EMMA #02074, ^45^) with mice expressing the CRE recombinase under the control of the *Villin* promoter. As controls we used littermate Hk2^fl/fl^ mice, referred to as WT mice. All mice were kept under a 12-h light cycle and fed gamma-irradiated diet *ad libitum*. Mice were killed by cervical dislocation prior to removing tissues for histological and molecular analyses. For basal phenotyping we used 9 to 11 and 86 to 92 weeks old mice. For DSS-induced colitis we used 10 to 14 weeks old male mice. Both genotypes were co-housed throughout the entire experiment. To induce colitis, mice received 1,5% (w/v) dextran sodium sulfate in autoclaved tap water. For the SCFA intervention mice were fed either a control diet, a butyrate-enriched diet or an acetate-containing diet for ten days prior to inducing colitis by administering DSS. The SCFA supplementation was continued throughout the entire experiment. The butyrate-enriched diet consisted of control feed (V1534, ssniff) supplemented with 10% (w/w) of the butyrate-polymer tributyrin (Sigma Aldrich), as this ensures the release of butyrate in the colon after metabolization by the intestinal microbiota instead of its absorption in the proximal small intestine ^46,47^. The acetate-enriched diet consisted of control feed but the drinking water was replaced with autoclaved tap water supplemented with 150 mM sodium acetate (Sigma Aldrich), a concentration used successfully in previous studies ^26,36^. Gnotobiotic experiments were performed in the animal facilities of either the Experimental Biomedicine (EBM) of Gothenburg or Hannover Medical School (MHH). All animal protocols were approved by the Gothenburg Animal Ethics Committee or by Lower Saxony State Office for Consumer Protection and Food Safety. Gnotobiotic mice were housed as described under standard procedures ^48,49^. Mice were kept under a 12-h light cycle and fed autoclaved chow diet *ad libitum* (Labdiet, St Louis, MO, USA). Monoassociated mice were generated by inoculating 12-week-old GF mice with 200 μl of overnight (stationary phase) *in vitro* cultures of *B. thetaiotaomicron*, *E. coli* or *B. longum* by oral gavage and the mice were sacrificed 14 days post colonization. ASF- and OMM-associated mice were generated by co-housing of weaned germ-free mice for four weeks with gnotobiotic donor animals colonized with either ASF or OMM and sacrificed at an age of 12 weeks.

### Bacteria and *in vitro* culture

*B. thetaiotaomicron* VPI-5482 (ATCC 29148, ^50^), *E. coli* W3110 ^51^ were kindly provided by Dr. Jeffrey Gordon (Edison Family Center for Genome Sciences and Systems Biology, Washington University School of Medicine, St. Louis, MO 63110, USA), while *B. longum* NCC 2705 was kindly provided by Dr. Stéphane Duboux, Nestlé Research Center, Lausanne, Switzerland. Liquid media (TYG, LB and MRS broth supplemented with 0.05% (w/v) cysteine, for *B. thetaiotaomicron*, *E. coli* and *B. longum*, respectively) were inoculated with single colonies of cultures on agar plates and were grown to the stationary phase overnight in an anaerobic jar at 37°C. ASF is a mix of eight bacteria: two Clostridia species ASF356 & ASF502, *Lactobacillus murinus* ASF361 and *spec*. ASF360, *Mucispirillum schaedleri* ASF457, *Eubacterium plexicaudatum* ASF492, *Parabacteroides spec*. ASF519 and an unknown Firmicutes bacterium ASF500 ^52^. ASF colonized mice were purchased from Taconic and inoculated transgenerally by co-housing. OMM (Oligo-Mouse-Microbiota) is a mix of 12 bacteria: *Bacterioides ceacimuris* I48*, Muribaculum intestinale* YL27*, Akkermansia municiphila* YL44*, Turicimonas muris* YL45*, Limosilactobacillus reuteri* I49*, Enterococcus faecalis* KB1*, Blautia coccoides* YL58*, Clostridium innocuum* I46*, Flavonifractor plautii* YL31*, Enterocloster clostridioforme* YL32*, Acutalibacter muris* KB18 *and Bifidobacterium animalis* YL2 ^21^. Bacteria of the OMM consortium were grown in single cultures under anaerobic conditions in Anaerobic Akkermansia Medium (AAM) as previously described ^21^.

### Isolation of primary cells and intestinal organoids

IECs were isolated from intestinal tissue using the Lamina Propria Dissociation Kit (Miltenyi BioTech, Bergisch Gladbach, Germany) according to the manufacturer’s protocol with minor deviations as described before ^53^. In brief, intestinal epithelial cells were isolated by disruption of the structural integrity of the epithelium using ethylenediaminetetraacetic acid (EDTA) and dithiothreitol (DTT). Purity of individual IEC fractions was analyzed by flow cytometry on a FACS Calibur flow cytometer (B&D, Heidelberg, Germany) with Cellquest analysis software from Becton Dickinson. We used the Anti-EpCam-PE (Clone: G8.8, Biolegend, San Diego, USA) antibody for analysis of IEC purity. FACS-data was analysed using Flowing Software (Perttu Terho, Turku Centre for Biotechnology, Finland). Intestinal organoids were generated following procedures described earlier by Sato et al., 2009 ^54^. Organoids were cultivated in ENR-conditioned medium supplemented with human recombinant EGF as described by Sato et al., 2011 ^55^.

### Generation and culture of HK2-deficient Caco-2 cells

A CRISPR plasmid targeting human *Hk2* was generated using the GeneArt^®^ CRISPR Nuclease Vector Kit from Thermo Fisher. Caco-2 cells were purchased from DSMZ (ACC-169) and transfected with the *Hk2* CRISPR plasmid using Lipofectamin reagent kit (Thermo Fisher Scientific). Positive clones were screened via Western blot to generate a monoclonal population termed Caco-2^ΔHk2^. Caco-2 cells that were also subjected to the CRISPR transfection and selection procedure, but which still showed a HK2 band as per western blot analysis were used as controls (Caco-2^WT^). Caco-2^WT^ and Caco-2^ΔHk2^ cells were cultured in MEM with 20% (v/v) FCS purchased from Gibco/Life Technologies. Cells were seeded with 70% confluency and 24 hours in advance of stimulation.

### RNA isolation and qPCR

Total RNA was extracted using the RNeasy Mini Kit (Qiagen) according to the manufacturer’s protocol. RNA concentration was measured using a NanoDrop ND-1000 spectrophotometer (PeqLab Biotechnologie). 1μg of total RNA was reverse-transcribed to cDNA using the Maxima H Minus First Strand cDNA Synthesis Kit (ThermoFisher Scientific). qPCR was carried out using SYBR Select Master Mix (Applied Biosystems) according to the manufacturer’s instructions on a Viia 7 Real-Time PCR System (ThermoFisher Scientific). Expression levels were normalized to *Actb* (β-actin or *Rpl32* (ribosomal protein L32). Primer sequences are listed in Extended data table 3.

### RNA sequencing

Total RNA was extracted as described above from isolated IECs from *Hk2*^Δ*IEC*^ and littermate control mice under untreated conditions and after administering DSS for 3 or 7 days. RNA libraries were prepared using TruSeq stranded mRNA Kit (Illumina) according to manufacturer’s instructions. All samples were sequenced using an Illumina NovaSeq 6000 sequencer (Illumina, San Diego,CA) with an average of 23 million paired-end reads (2x 50 bp) at IKMB NGS core facilities. The RNA-seq data was processed using an inhouse pipeline (https://github.com/nf-core/rnaseq). Briefly, adapters and low-quality bases from the RNA-seq reads were removed using Trim Galore (version 0.4.4). The filtered reads were mapped to the mouse genome (GRCm38) using STAR aligner (version 2.5.2b). Expression counts of the transcripts were estimated using featureCounts (version 1.5.2) and then normalized across samples using the DESeq normalization method. DEseq2 ^56^ was used to determine differentially expressed genes. Genes were considered as significant differentially expressed if the adjusted *p-value* (Benjamini-Hochberg (BH) multiple test correction method) was less than 0.05. Gene Ontology (GO) enrichment analysis was conducted using the Bioconductor package topGO (version 2.32.0) and a Fisher.elim p-value (weight algorithm) of 0.05 was used as significance threshold.

### Microbiota analysis

MiSeq 16S amplicon sequence data was analyzed using MacQIIME v1.9.1 (http://www.wernerlab.org/software/macqiime, ^57^) as described previously ^58,59^. Briefly, all sequencing reads were trimmed keeping only nucleotides with a Phred quality score of at least 20, then paired-end assembled and mapped onto the different samples using the barcode information. Sequences were assigned to operational taxonomic units (OTUs) using uclust and the greengenes reference database (gg_13_8 release) with 97% identity ^60,61^. Representative OTUs were picked and taxonomy assigned using uclust and the greengenes database. Quality filtering was performed by removing chimeric sequences using ChimeraSlayer ^62^ and by removing singletons and sequences that failed to align with PyNAST ^63^. The reference phylogenetic tree was constructed using FastTree 2 ^64^. All samples within a single analysis were normalized by rarefaction to the minimum shared read count to account for differential sequencing depth among samples (10,174 sequences per sample). Relative abundance was calculated by dividing the number of reads for an OTU by the total number of sequences in the sample. Alpha diversity measures were computed and beta diversity was calculated using Unweighted Unifrac and visualized by principal coordinate analysis. Significance of differences in abundances of various taxonomic units between *Hk2*^Δ*IEC*^ and littermate control mice was calculated using t-test and p values were adjusted for multiple testing using FDR correction (q-value).

### Western blot analyses

Caco-2 cells were lysed using RIPA buffer. Organoids were lysed using SDS-based DLB buffer + 1% Halt Protease inhibitor cocktail (Thermo Fisher Scientific). Lysates were heated to 95°C for 5 min centrifuged at 16,000 g for 15 min at 4°C to remove cell remnants. Protein concentrations were measured by DC Protein Assay (BioRad) according to the manufacturers protocol. Equal amounts of lysates containing Laemmli buffer were heated at 95°C and electrophoresed on 12% polyacrylamide gels under standard SDS-PAGE conditions before being transferred onto a polyvinylidene fluoride membranes (GE Healthcare). Protein loaded membranes were blocked with 5% (w/v) non-fat dry milk or bovine serum albumin (BSA) in Tris-buffered saline (TBS) supplemented with 0,1% (v/v) Tween 20 for 1 hour, incubated with primary antibody (mouse anti-HK2, Novus Biologicals, #NBP2-02272; rabbit anti-PARP1/cPARP1, Cell Signaling Technology, #9542; rabbit anti-Caspase3/cCaspase3, Cell Signaling Technology, #9662, mouse anti-betaActin, Abcam, #ab20272) overnight, washed three times with TBS-Tween-20 and then incubated with the secondary horseradish peroxidase (HRP)-conjugated antibody for 1 hour at room temperature. Proteins were detected using the Pierce ECL and ECL Plus Substrate Kits (ThermoFisher).

### Histology and immunostaining

Tissue specimen were fixed in 10% formalin solution over night at 4°C and then embedded in paraffin. 5 μm thick sections were cut and stained with hematoxylin and eosin (H&E) or subjected to immunostaining using the Vectastain ABC kit (Vector Laboratories) including antigen retrieval in boiling citrate buffer. Primary antibodies were incubated overnight. For immunostaining of HK2 we used a 1:1000 diluted antibody (Novus Biologicals, #NBP2-02272). For immunostaining of Ki67 we used a 1:500 diluted mouse anti-Ki67 antibody (BD Biosciences, cat.no. 556003). The TUNEL assay was performed using the ApopTag Plus Peroxidase In Situ Apoptosis Detection Kit (Merck Millipore) according to the manufacturer’s instructions. Slides were visualized using a Zeiss Imager Z1 microscope (Zeiss) and pictures were taken using ZEN pro (Zeiss) software.

### Seahorse analysis

To perform real-time ECAR and OCR-analyses, Caco-2 cells were analyzed using the Seahorse XF24 Analyzer from Agilent Technologies by Mito Stress Test Kit or Glycolysis Stress Test Kit according to the manufacturer’s instructions. 4×10^4^ cells were used in each assay with n=9 replicates in three independent experiments.

### Metabolic modelling of SCFA production by OMM bacteria

Metabolic networks were reconstructed using gapseq ^65^ based on the genomes of OMM bacteria, which were downloaded from NCBI bioproject PRJNA289613, and the Anaerobic Akkermansia Medium ^21^. The formation of fermentation products was determined via an extended flux balance analysis that minimizes the total flux through all reactions as a proxy for the parsimonious enzyme usage ^66^. The variability of predictions was taken into account by randomly sampling the space of alternative optimal solutions via the function ACHR from the R package sybilcycleFreeFlux (W=5000, nPoints=10000) ^67^ to derive the distribution of fermentation products for each bacteria. All predicted fermentation products were then used as explanatory variables to estimate the *Hk2* expression via linear regression.

### Quantification of SCFAs

OMM bacteria were grown as described and 1 mL of each bacterial culture supernatant was homogenized using NucleoSpin Bead Tubes (Macherey-Nagel, Düren, Germany) and a Precellys Evolution Homogenizer (Bertin Corp., Rockville, Maryland, USA). Homogenates were cleared by centrifugation for 10 min at 21,000 × g and 4°C. SCFA standards including acetic, propionic and butyric acid (purchased from Sigma Aldrich, St Louis, MO, USA) were prepared in methanol to a concentration of 100 ppm. Derivatization of SCFAs was performed with 3-nitrophenylhydrazone as described ^68^. SCFA concentrations in the samples were then measured via Ultra-High Performance Liquid Chromatography (Acquity UPLC, Waters, Milford, MA, USA) coupled to Mass Spectrometry (amaZon ETD IonTrap, Bruker Daltonics GmbH, Bremen, Germany). Sample separation was performed using a C8 column and solvent system consisting of ammonium acetate (5 mM, Sigma Aldrich, St Louis, MO, USA) combined with acetic acid (0.1%, pH 4.2, Biosolve, Valkenswaard, Netherlands) in water or acetonitrile (LC-MS CHROMASOLV, FLUKA, Sigma Aldrich, St Louis, MO, USA). Mass spectrometry analysis was performed in negative electrospray ionization mode. Peak areas and concentrations of SCFAs in the bacterial supernatants were calculated using QuantAnalysis (Bruker, Daltonics, Bremen, Germany) software.

### Statistical analysis

Biostatistical analyses were performed using GraphPad Prism (version 8) software (GraphPad, Inc, La Jolla, CA), MacQIIME v1.9.2 or R (v 3.2.5). Specific comparisons and analyses are described in the individual method sections. Differences between the groups were considered significant at P < 0.05 and the data are presented as means ± SEM.

### Data Availability Statement

All data is either included in this manuscript or deposited on public databases. The RNA sequencing data has been deposited at NCBI’s Sequence Read Archive under the accession number GSE158026. The 16S amplicon sequencing data are accessible through the European Nucleotide Archive (ENA, https://www.ebi.ac.uk/ena) under the study accession number PRJEB40281. Additional data that support the findings of this study are available from the corresponding author upon reasonable request.

### Code Availability Statement

All codes used to generate the bioinformatic analyses are available from the corresponding author upon reasonable request.

## Acknowledgments

The authors thank Karina Greve, Dorina Oelsner, Melanie Vollstedt, Maren Reffelmann, Melanie Nebendahl, Sabine Kock and Stefanie Baumgarten for excellent technical assistance. We also thank Dr. Filipe de Vadder (Lyon University, France) for critical comments that greatly improved the manuscript.

This work was supported by the German Research Foundation (DFG) through the individual grant SO1141/10-1, the Research Unit FOR5042 “miTarget - The Microbiome as a Target in Inflammatory Bowel Diseases” (project P5), the Collaborative Research Centre CRC1182 “Origin and Function of Metaorganisms” (project C2), the Excellence Clusters EXS2167 “Precision Medicine in Chronic Inflammation” and EXC306 “Inflammation at Interfaces” and via the SH Excellence Chair Program to Philip Rosenstiel.

## Author contributions

JH, FH, LS, PR and FS designed research. JH, FH, LS, NS and FS managed the conventional mouse colony, monitored the mice, performed animal experiments and collected samples. MB, VT, FB maintained the gnotobiotic mouse facilities, performed the colonizations of GF mice with single bacteria and minimal consortia and provided samples. JH, FH, LS, AW, KS, NS, KK, DP, SK and PR conducted the wet lab experiments. JH, FH, LS, NM, KS, NS, KK, RH, SK, FB, PR and FS analyzed and interpreted the data. JH, NM, AW, JZ, RH and FS performed the bioinformatics analyses. JH, FH, LS and FS prepared the figures. PR and FS obtained funding. JH, FH, LS, PR and FS co-wrote the manuscript with critical input from all authors.

## Competing interest declaration

The authors have no competing interests to declare. All authors have read and approved the manuscript and agree with its submission. The manuscript has not been previously published and is not currently under consideration by another journal.

## Additional information

### Abbreviations

ASF: altered Schaedler Flora
BSA: bovine serum albumin
Caco-2: colon car-cinoma cell line 2
CD: Crohn’s disease
CR: conventionally raised mice
DAI: disease activity index
DSS: dextran sodium sulfate
ECAR: extracellular acidification rate
EWAT: epididymal white adipose tissue
FCCP: cyanide-4-(trifluo-romethoxy)-phenylhydrazone
FFAR: free fatty-acid receptor
GF: germ-free
GO: gene ontology
GPR: G protein-coupled receptor
H&E: hematoxylin and eosin
HDAC: histone deacetylases
HK: hexokinase
HRP: horseradish peroxidase
IBD: inflammatory bowel disease
IEC: intestinal epithelial cell
MPTP: mitochondrial permeability transition pore
OCR: oxygen consumption rate
OMM: oligo Mouse Microbiota
PARP1: Poly(ADP-Ribose)-Polymerase 1
PPIF: peptidyl-prolyl cis-trans isomerase
qPCR: quantitative real-time polymerase chain reaction
SCFA: short-chain fatty acid
SPF: specific pathogen free
TBS: Tris-buffered saline
TNF: tumor necrosis factor
TUNEL: terminal deoxynucleotidyl transferase dUTP nick end labelling
UC: ulcerative colitis
WT: wild-type

## Extended data figure/table legends

**Extended data figure 1:**
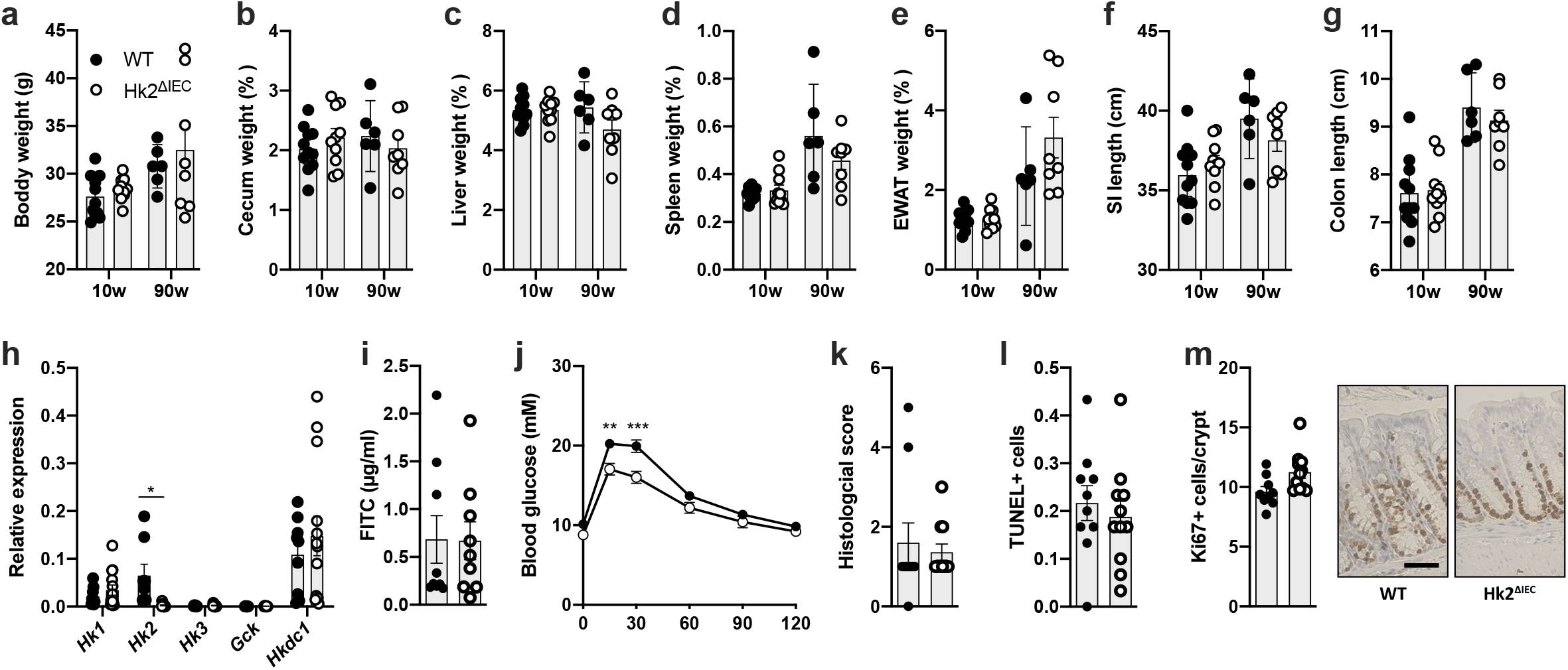
*Hk2*^ΔIEC^ mice do not display an evident immunological or metabolic phenotype under baseline conditions. **a-g)** 10- or 90-week-old WT and *Hk2*^ΔIEC^ mice were sacrificed and organ measures taken. **a)** Body weight. **b)** Cecum weight. **c)** Liver weight. **d)** Spleen weight. **e)** Epididymal white adipose tissue (EWAT) weight. **f)** Small intestine length. **g)** Colon length. **h)** Relative expression of all hexokinase family members in unfractionated colon tissue of 10-week-old WT and *Hk2*^ΔIEC^ mice as per qPCR. **i)** Intestinal permeability as measured by fluorescein isothiocyanate (FITC) levels in blood of 10-week-old WT and *Hk2*^ΔIEC^ mice 1h after oral gavage. **j)** Blood glucose levels during oral glucose tolerance test in 10-week-old WT and *Hk2*^Δ*IEC*^ mice. **k)** Colonic histological score of 10-week-old WT and *Hk2^ΔIEC^* mice. **l)** TUNEL-positive cells per colon crypt of 10-week-old WT and *Hk2*^ΔIEC^ mice. **m)** Ki-67-positive cells per colon crypt of 10-week-old WT and *Hk2*^ΔIEC^ mice including representative images. The scale bar represents 50 μm.

**Extended data figure 2:**
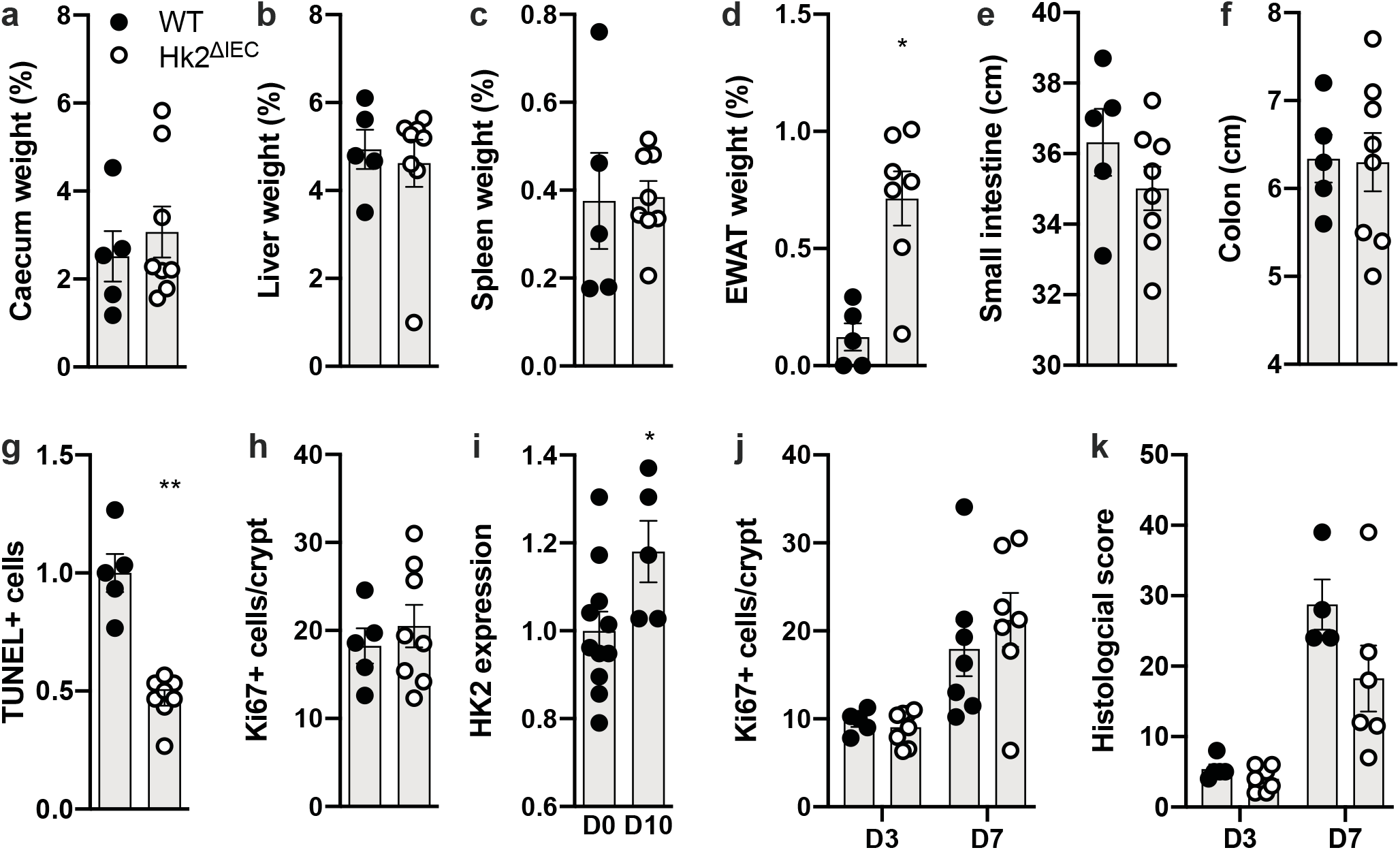
Phenotyping of WT and *Hk2*^ΔIEC^ mice during DSS-induced colitis. **a-h)** Organ and histological data from WT and *Hk2*^ΔIEC^ mice sacrificed at the end (day 10) of DSS colitis. **a)** Cecum weight. **b)** Liver weight. **c)** Spleen weight. **d)** Epididymal white adipose tissue (EWAT) weight. **e)** Small intestine length. **f)** Colon length. **g)** TUNEL-positive cells per colon crypt. **h)** Ki-67-positive cells per colon crypt. **i)** HK2 protein levels in colon epithelium of WT mice at baseline (day 0) and at day 10 during DSS colitis as determined per immunohistochemistry. **j)** Ki-67-positive cells per colon crypt and **k)** histological score of WT and *Hk2*^ΔIEC^ mice at day 3 and 7 during DSS colitis.

**Extended data figure 3:**
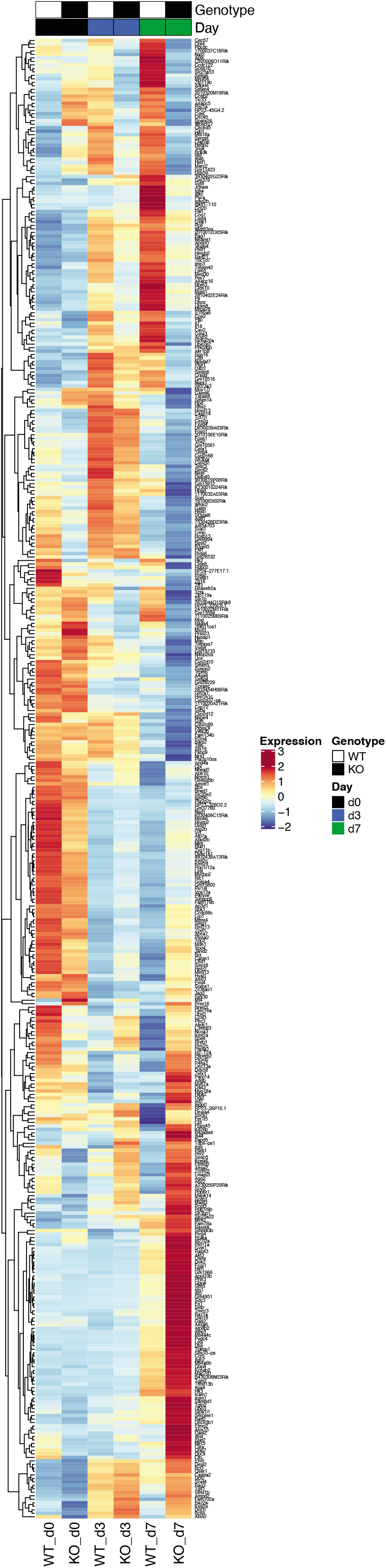
Heatmap of genes differentially expressed in colon of WT and *Hk2*^ΔIEC^ mice at days 0, 3 and 7 during DSS colitis. WT and *Hk2*^ΔIEC^ mice were given 2% DSS in drinking water and analyzed on day 0, 3 and 7 of treatment. RNA was isolated from unfractionated colon and sequenced to identify transcripts with differential expression dependent on the loss of epithelial HK2 during the onset of inflammation.

**Extended data Figure 4:**
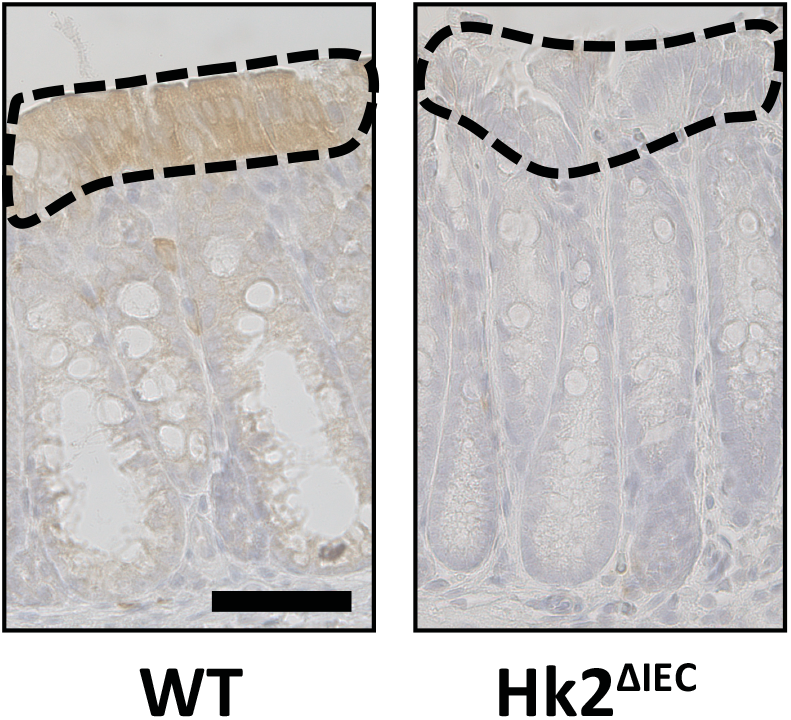
Expression pattern of the HK2 protein in the colonic epithelium of untreated 10-week-old WT and *Hk2*^ΔIEC^ mice. The scale bar represents 50 μm. The dotted line indicates the tip epithelium area used for quantification of HK2 protein expression. Note the dominant HK2 expression in the colonic tip epithelium.

**Extended data Figure 5:**
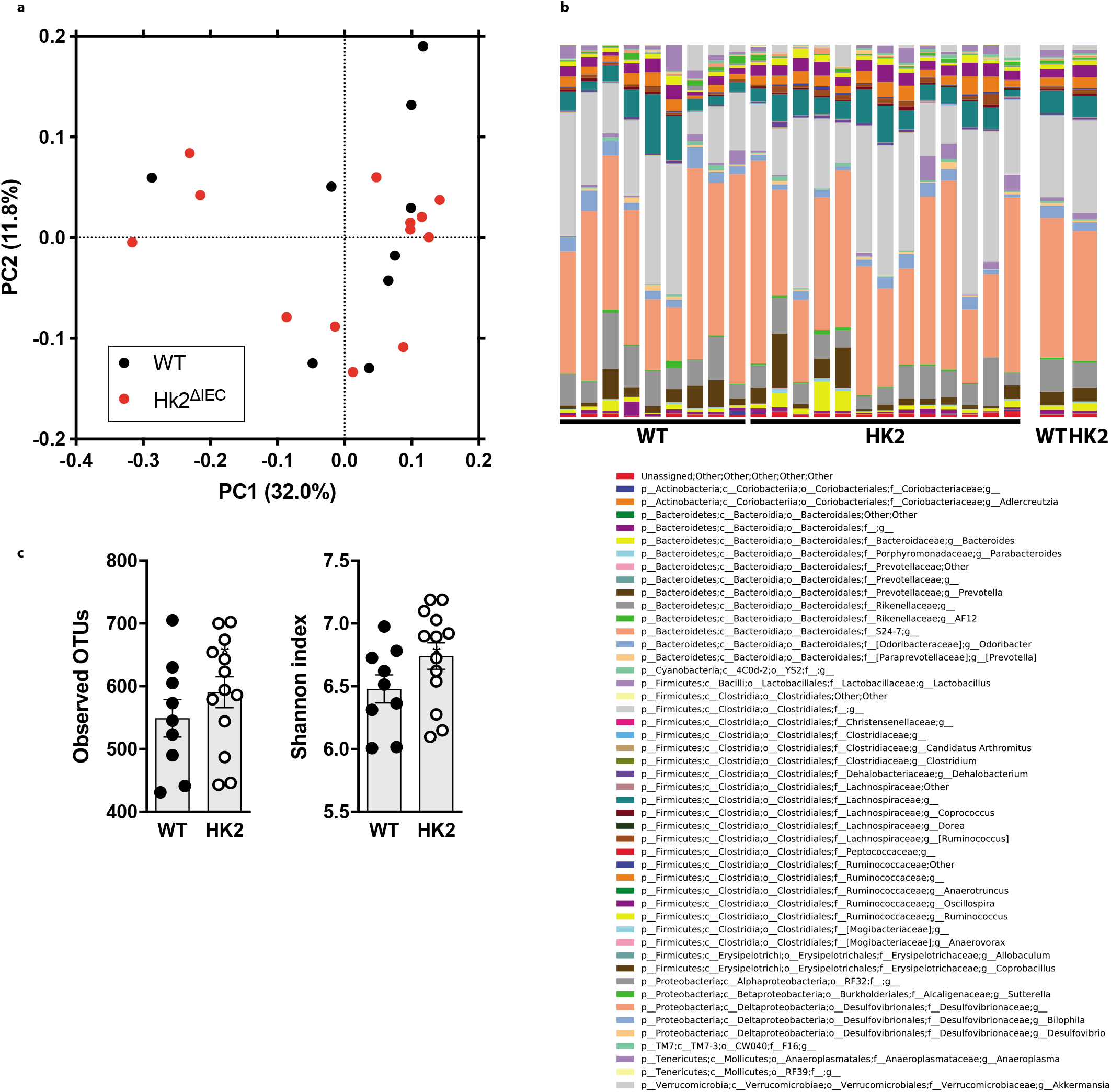
Ablation of HK2 in IECs does not alter the composition of the intestinal microbiota. **a)** Principal coordinate analysis of fecal microbiota from WT and *Hk2*^ΔIEC^ mice. **b)** Taxonomic overview on genus level. **c)** Alpha diversity (the variation of microorganisms in a single sample).

**Extended data Figure 6:**
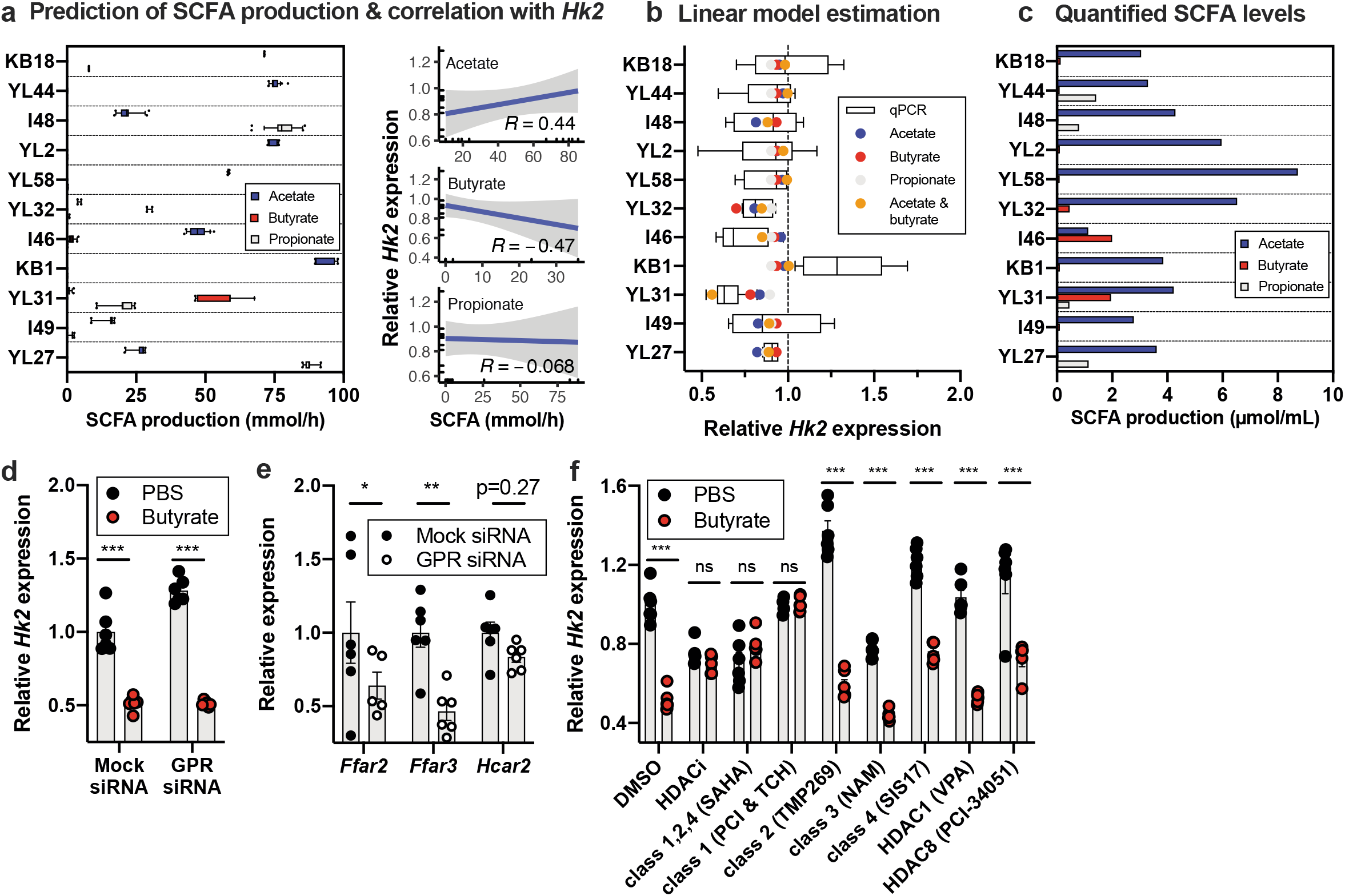
Role of SCFAs and HDACs in butyrate-mediated repression of *Hk2* expression. **a)** Metabolic modeling of the production of acetate, butyrate and propionate by the individual OMM bacteria based on their published genome information and the used *in-vitro* growth conditions, and correlation of metabolite production with *Hk2* expression. R denotes the Pearson correlation coefficient. **b)** Prediction of linear models incorporating the acetate, butyrate and propionate levels to explain the observed changes in *Hk2* expression upon stimulation of Caco-2 cells with the bacterial culture supernatants (shown in Fig. 3d). **c)** SCFA levels quantified in culture supernatants of single OMM bacteria, which were used to stimulate Caco-2 cells (shown in Fig. 3d). **d)** Relative *Hk2* expression in Caco-2 cells after transfection with a mix of siRNA targeting the three G protein-coupled receptors (GPR) GPR41 (*Ffar3*), GPR43 (*Ffar2*) and GPR109a (*Hca2*) and subsequent stimulation with butyrate. **e)** Expression of *Ffar3, Ffar2* and *Hcar2* in Caco-2 cells after transfection with the siRNA targeting these three GPRs to test for successful knockdown. **f)** Relative *Hk2* expression in Caco-2 cells first treated with general or specific HDAC inhibitors and then incubated with butyrate. HDACi refers to a mixture of SAHA and NAM and was used to inhibit all HDAC classes. A mixture of PCI-34051 and TC-H106 were used to inhibit class I HDACs. TMP269, NAM (nicotinamide) and SIS17 were used to inhibit class II, III and IV, respectively. Valproic acid (VPA) and PCI-34051 were used to inhibit only HDAC1 and HDAC8, respectively. All expression values were normalized to the mean of DMSO-PBS.

**Extended data figure 7:**
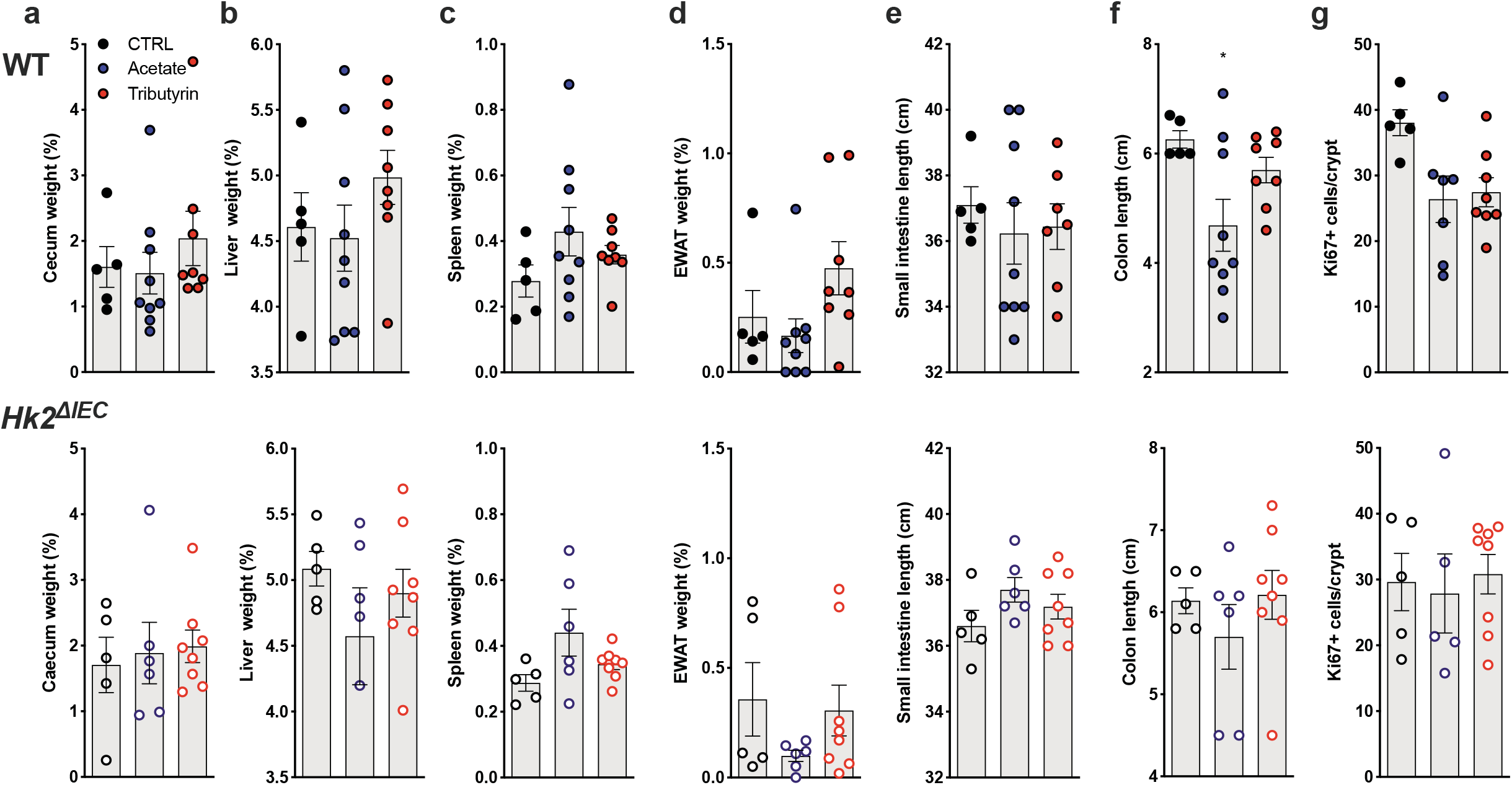
Organ measures and histological data of WT and *Hk2*^ΔIEC^ mice at the end of the dietary SCFA supplementation and DSS-induced colitis experiment. **a)** Cecum weight. **b)** Liver weight. **c)** Spleen weight. **d)** Epididymal white adipose tissue (EWAT) weight. **e)** Small intestine length. **f)** Colon length. **g)** Ki-67-positive cells per colon crypt.

**Extended data table 1: Complete normalized read counts and statistics from RNA sequencing of of IECs isolated from colon of WT and *Hk2*^ΔIEC^ mice at day 0 (baseline) and days 3 and 7 of DSS colitis.**

**Extended data table 2: Statistics of linear modelling of *Hk2* expression induced by metabolites produced by single OMM bacteria grown *in vitro*.** AIC = Akaike Information Criteria. Sigma = standard deviation of the residuals. R^2^ = coefficient of determination, referring to the proportion of variance in the dependent variable that is predictable from independent variables. R = Pearson correlation coefficient of predicted metabolite production with *Hk2* expression data.

**Extended data table 3: List of primers used in this study.**

## Notes

### Competing Interest Statement

The authors have declared no competing interest.

